# The Ensembl Variant Effect Predictor

**DOI:** 10.1101/042374

**Authors:** William McLaren, Laurent Gil, Sarah E Hunt, Harpreet Singh Riat, Graham R. S. Ritchie, Anja Thormann, Paul Flicek, Fiona Cunningham

**Affiliations:** European Molecular Biology Laboratory, European Bioinformatics Institute, Wellcome Genome Campus, Hinxton, Cambridge, CB10 1SD, United Kingdom.

**Keywords:** variant annotation, NGS, genome, SNP

## Abstract

The Ensembl Variant Effect Predictor (VEP) is a powerful toolset for the analysis, annotation and prioritization of genomic variants, including in non-coding regions.

The VEP accurately predicts the effects of sequence variants on transcripts, protein products, regulatory regions and binding motifs by leveraging the high quality, broad scope, and integrated nature of the Ensembl databases. In addition, it enables comparison with a large collection of existing publicly available variation data within Ensembl to provide insights into population and ancestral genetics, phenotypes and disease.

The VEP is open source and free to use. It is available via a simple web interface (http://www.ensembl.org/vep), a powerful downloadable package, and both Ensembl’s Perl and REST application program interface (API) services.

## INTRODUCTION

Analysis of variant data resulting from genome or exome sequencing is fundamental for progress in biology, from basic research to translational genomics in the clinic. It is key for investigating function and for progressing from a system of medical care based on standardized treatment to one targeted to the individual patient

For sufferers of common or rare disease, the potential benefits of variant analysis include improving patient care, surveillance and treatment outcomes. In cancer, there have already been numerous successes using data from genetic tests. For example, patients testing positive for the inheritance of BRCA mutations have the option of selective preventative surgery; lung cancer patients showing EGFR gene mutations, or triple negative breast cancer patients can have their drug prescriptions tailored to improve success (Eisenstein 2014; Weil and Chen 2011).

Rare diseases can individually be difficult to diagnose due to the low incidence and the incomplete penetrance of implicated alleles. However, variant analysis of whole genome sequencing (WGS) or whole exome sequencing (WES) data can lead to the discovery of underlying genetic mutations (The Deciphering Developmental Disorders Study 2015). Identifying an associated mutation is advantageous for researching treatment options and for future drug discovery. Meanwhile even the immediate benefit of diagnosis may result in a more accurate prognosis and remove the burden of additional medical investigations.

The most common non-infectious diseases worldwide are cardiovascular disease, cancer and diabetes (World Health Organisation 2015). Despite many array-based genome-wide association studies searching for risk loci, only a relatively small heritable component in these conditions has been elucidated (Visscher et al. 2012). WGS in large numbers of samples is required to yield enough statistical power to detect rare variants with potential phenotypic or disease associations (Pierre and Génin 2014; Zuk et al. 2014). WGS studies will also detect variants in regulatory and non-coding regions of the genome, which are thought to comprise the majority of trait-associated variants (Hindorff et al. 2009) and play a role in cancer (Puente et al. 2015).

The potential of large-scale sequencing and variant analysis is revolutionary. Recognising this value, major population sequencing initiatives have been launched in Iceland (Gudbjartsson et al. 2015), the UK (100,000 Genomes Project 2014) and the USA (Precision Medicine Initiative, Collins and Varmus 2015). In other species, efforts such as Genome 10K (Koepfli et al. 2015), the 1001 Arabidopsis genomes (Cao et al. 2011) and 1000 bull genome project (Daetwyler et al. 2014) have similar goals, but operate under different funding models, often with less support than the *Homo sapiens* focused projects.

Ongoing improvements in DNA sequencing technology, and a current cost around $1000 per human genome, have resulted in high volumes of genome, exome and subsequent variant data requiring interpretation. Meanwhile the cost of the analysis to determine functional consequences remains substantially higher due to the difficulty of variant interpretation. For example, a typical diploid human genome has around 3.5 million single nucleotide variants (SNVs) and 1,000 copy number variants (Gonzaga-Jauregui et al. 2012) with respect to the genome reference sequence. Around 20,000 – 25,000 of these variants are protein coding, 10,000 change an amino acid but only 50 – 100 of these are protein truncating or loss of function variants (Gonzaga-Jauregui et al. 2012). Manual review of large numbers of variants is impractical and costly, and there are additional difficulties, such as a lack of functional annotation or the interpretation of multiple variants within a haplotype.

Variant interpretation often considers the impact of a variant on a transcript or protein. It is therefore dependent on transcript annotation and localizing variants to protein coding or non-coding regions. There are two major sources of *Homo sapiens* annotation: GENCODE (Harrow et al. 2012) and Reference Sequence (RefSeq, Pruitt et al. 2014) at the National Center for Biotechnology Information (NCBI). Both sets of transcript annotation are subject to version changes and updates that can modify variant reporting and interpretation. For data reproducibility, transcript isoforms and transcript versions must be rigorously tracked, although in some cases even including the version is not sufficient to avoid all potential misinterpretations (Dalgleish et al. 2010).There are differences in how the transcript sets are produced: GENCODE annotation is genome-based while RefSeq transcripts are independent of the reference genome. Although RefSeq transcripts may correct for errors in the reference assembly and provide transcripts with improved biological representation (such as for the genes ABO, ACTN3 and ALMS1 in the GRCh37 reference), differences between a genome and a transcript set can cause confusion and errors when reporting variants at the cDNA and genomic level (e.g. these descriptions refer to the same variant: NM_000059.3:c.7397C>T, NC_000013.11:g.32355250T=). GENCODE’s aim is to create a comprehensive transcript set to represent expression of each isoform across any tissue and stage of development, and as a result there are on average nearly four transcripts per protein-coding gene (see Figure 1). Most genes, therefore, have several annotations for a given variant due to multiple transcript isoforms. This number will increase as more experimental data accumulates. Finally, in loci where the reference genome has several alternative haplotype representations (‘ALTs’) (Church et al. 2015), variants may have different interpretations with respect to different to ALTs. For example rs150580082 has mappings to multiple ALTs, where in only some of these it introduces a stop codon. In this case considering the primary assembly mapping alone will give misleading results.

**Figure 1:**
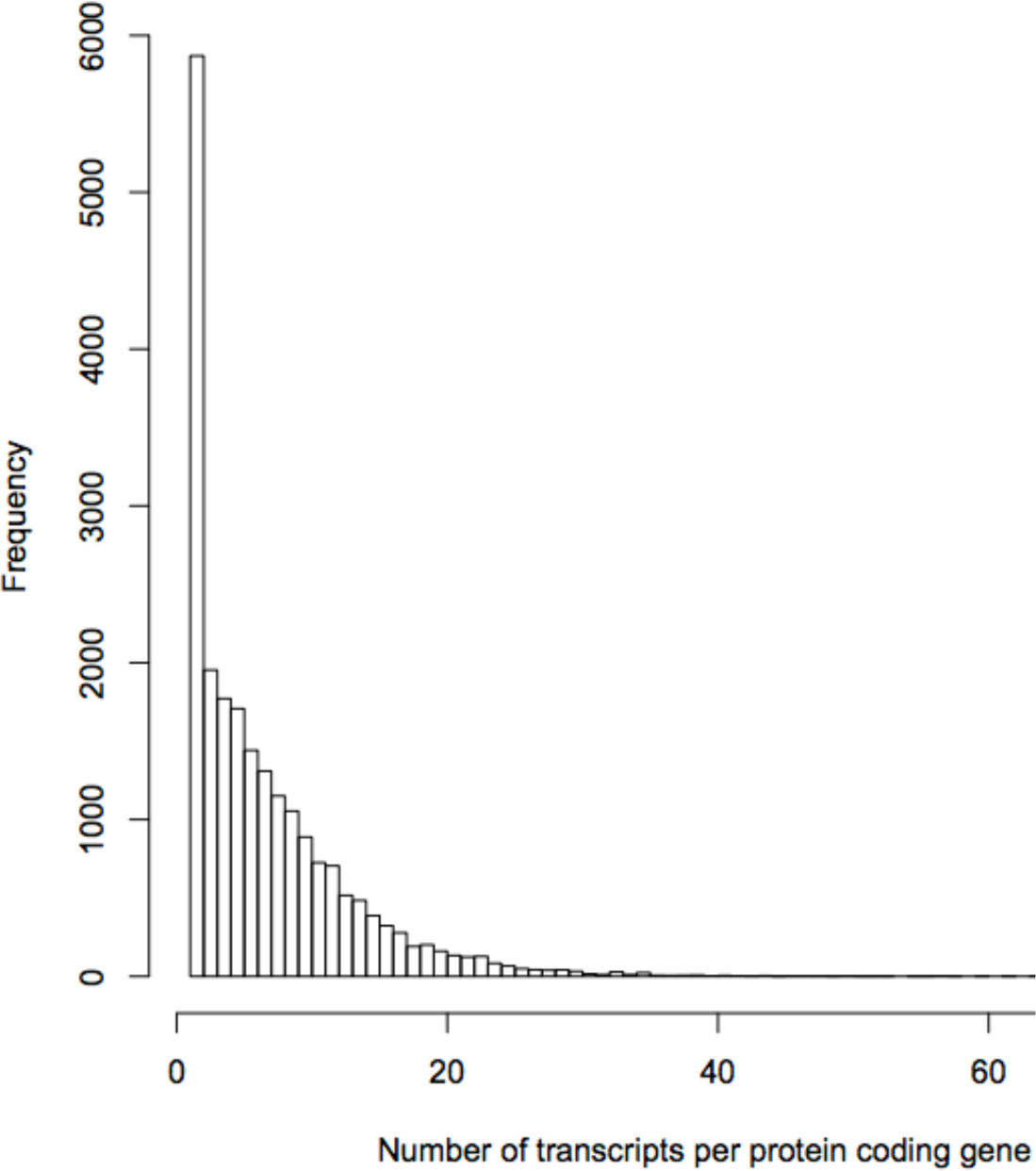
Each locus has the potential to have many transcript isoforms. For example in Ensembl release 79, there are 21,885 protein-coding genes that have an average of nearly four protein-coding transcripts. The maximum number of protein-coding transcripts, however, is 61 for the G protein-coupled receptor 56 gene (GPR56). For any changes to a transcript, GENCODE produces a new transcript version. Where there are many transcripts, or several versions of transcripts, choosing the correct transcript isoform and version for consistent variant annotation is challenging.

Variant reporting using Human Genome Variation Society (HGVS) nomenclature is also based on transcripts or proteins. Therefore the difficulties with transcript annotation described above may cause confusion and ambiguities when using HGVS nomenclature. Many possible annotations exist for variants in genes with multiple transcript isoforms. For example, rs121908462 is a pathogenic variant associated with Polymicrogyria that falls in ADGRG1, an adhesion G protein-coupled receptor G1. This variant has 126 HGVS descriptions in Ensembl (Cunningham et al. 2015) (and even more valid HGVS descriptions exist), as it overlaps 75 transcripts, and another 103 different descriptions in dbSNP. Multiple transcripts per locus result in greater numbers of annotations. These require filtering in a consistent manner, which increases the instability and complexity of variant interpretation.

Given these analysis challenges and the increasing volume of sequencing data being produced, there is a need for a robust computational tool to aid prioritization of variants across transcripts and manage the complexities of variant analysis. To facilitate this, we developed the Ensembl Variant Effect Predictor (VEP) (www.ensembl.org/vep), which differs significantly from other tools (Pabinger et al. 2014) (see Table 1 and discussion below) and from the previously published Ensembl SNP Effect Predictor (McLaren et al. 2010). The VEP is a software suite that performs annotation and analysis of most types of genomic variation in coding and non-coding regions of the genome. From disease investigation to population studies, it is a critical tool to annotate variants and prioritize a subset for further analysis.

**Table 1:**
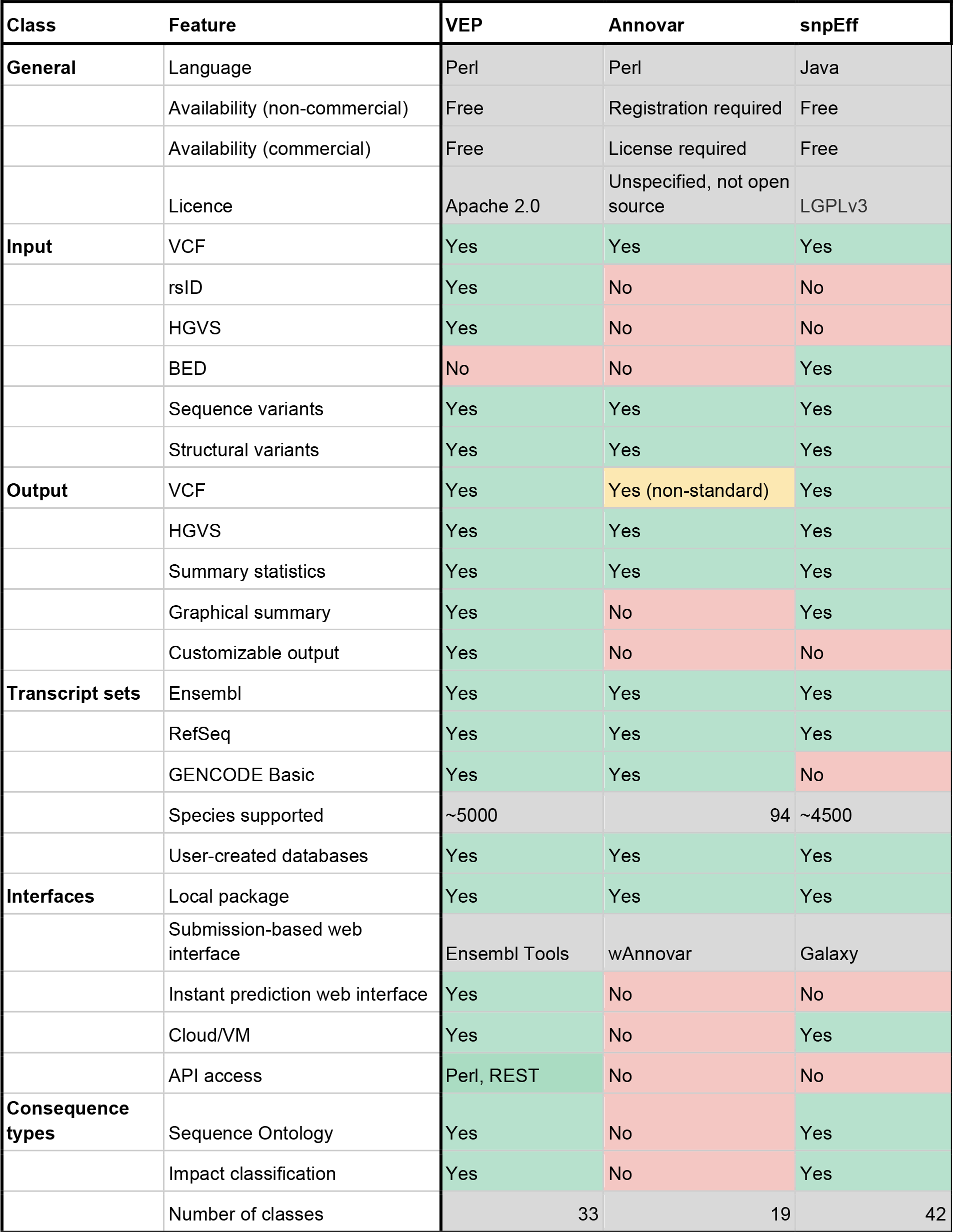
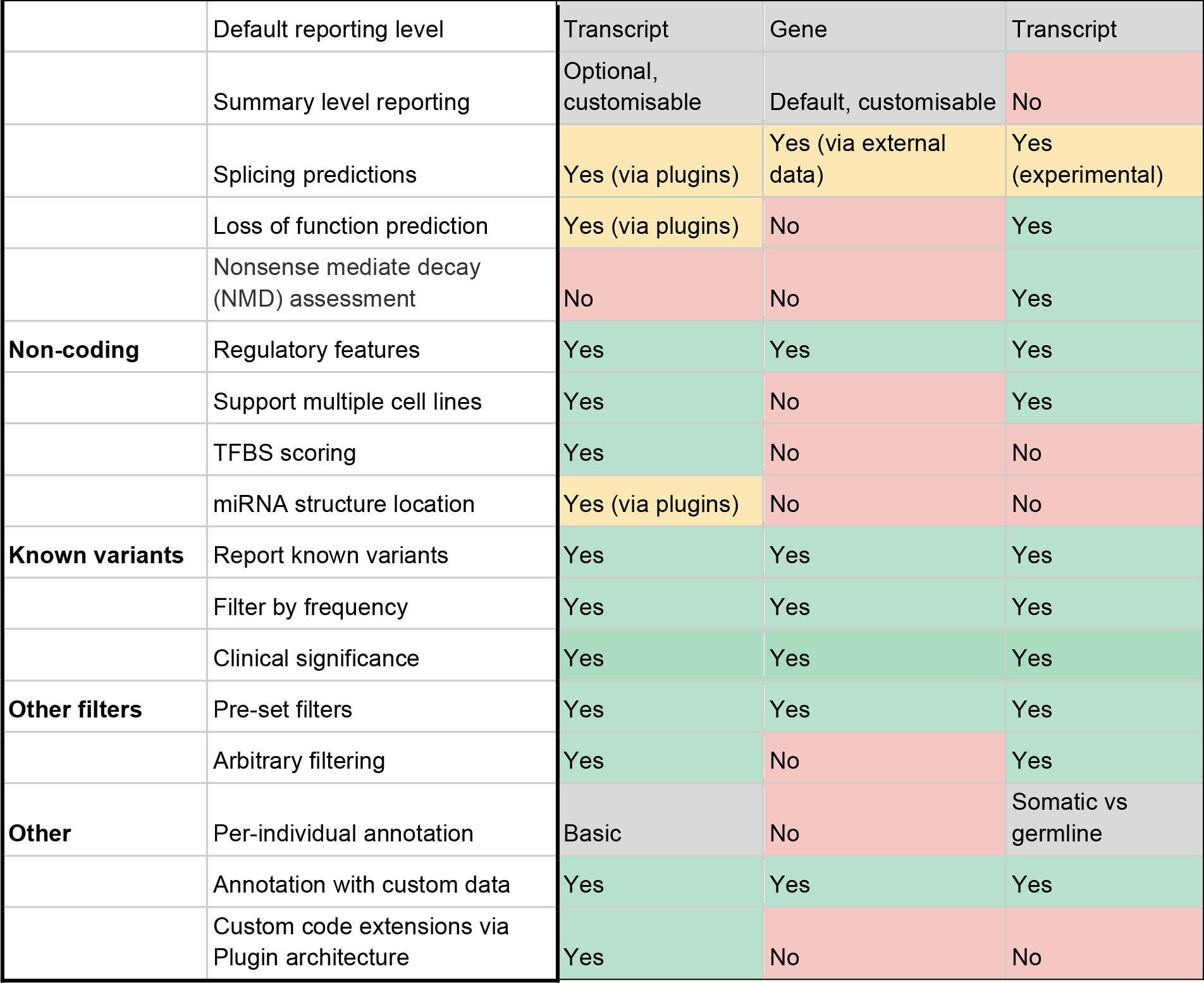
Comparison of features of VEP with Annovar (Wang et al. 2010) and snpEff (Cingolani et al. 2012).

| Class | Feature | VEP | Annovar | snpEff |
| --- | --- | --- | --- | --- |
| **General** | Language | Perl | Perl | Java |
|  | Availability (non-commercial) | Free | Registration required | Free |
|  | Availability (commercial) | Free | License required | Free |
|  | Licence | Apache 2.0 | Unspecified, not open source | LGPLv3 |
| **Input** | VCF | Yes | Yes | Yes |
|  | rsID | Yes | No | No |
|  | HGVS | Yes | No | No |
|  | BED | No | No | Yes |
|  | Sequence variants | Yes | Yes | Yes |
|  | Structural variants | Yes | Yes | Yes |
| **Output** | VCF | Yes | Yes (non-standard) | Yes |
|  | HGVS | Yes | Yes | Yes |
|  | Summary statistics | Yes | Yes | Yes |
|  | Graphical summary | Yes | No | Yes |
|  | Customizable output | Yes | No | No |
| **Transcript sets** | Ensembl | Yes | Yes | Yes |
|  | RefSeq | Yes | Yes | Yes |
|  | GENCODE Basic | Yes | Yes | No |
|  | Species supported | ~5000 | 94 | ~4500 |
|  | User-created databases | Yes | Yes | Yes |
| **Interfaces** | Local package | Yes | Yes | Yes |
|  | Submission-based web interface | Ensembl Tools | wAnnovar | Galaxy |
|  | Instant prediction web interface | Yes | No | No |
|  | Cloud/VM | Yes | No | Yes |
|  | API access | Perl, REST | No | No |
| **Consequence types** | Sequence Ontology | Yes | No | Yes |
|  | Impact classification | Yes | No | Yes |
|  | Number of classes | 33 | 19 | 42 |
|  | Default reporting level | Transcript | Gene | Transcript |
|  | Summary level reporting | Optional, customisable | Default, customisable | No |
|  | Splicing predictions | Yes (via plugins) | Yes (via external data) | Yes (experimental) |
|  | Loss of function prediction | Yes (via plugins) | No | Yes |
|  | Nonsense mediate decay (NMD) assessment | No | No | Yes |
| **Non-coding** | Regulatory features | Yes | Yes | Yes |
|  | Support multiple cell lines | Yes | No | Yes |
|  | TFBS scoring | Yes | No | No |
|  | miRNA structure location | Yes (via plugins) | No | No |
| **Known variants** | Report known variants | Yes | Yes | Yes |
|  | Filter by frequency | Yes | Yes | Yes |
|  | Clinical significance | Yes | Yes | Yes |
| **Other filters** | Pre-set filters | Yes | Yes | Yes |
|  | Arbitrary filtering | Yes | No | Yes |
| **Other** | Per-individual annotation | Basic | No | Somatic vs germline |
|  | Annotation with custom data | Yes | Yes | Yes |
|  | Custom code extensions via Plugin architecture | Yes | No | No |

The VEP has been used for analysis of traits in farm animals (Godoy et al. 2015; Höglund et al. 2014), for patient diagnosis in the clinic and for research on GWAS studies (Leslie et al.; Hou and Zhao 2013; International Multiple Sclerosis Genetics Consortium 2013; Saunders et al. 2012; Wright et al. 2015). It has been used for analysis in numerous large scale projects including the 1000 Genomes (McVean et al. 2012) and Exome Aggregation Consortium (ExAC) (http://exac.broadinstitute.org). It is a flexible tool of value to any project requiring detailed annotation of sequence variants.

## RESULTS

The VEP annotates two broad categories of genomic variant: (1) sequence variants with specific and well-defined changes (including SNVs, insertions, deletions, multiple base pair substitutions, microsatellites and tandem repeats); and (2) larger structural variants (greater than fifty nucleotides in length) including those with changes in copy number or insertions and deletions of DNA. For all input variants, the VEP returns detailed annotation for effects on transcripts, proteins and regulatory regions. For known or overlapping variants, allele frequencies and disease or phenotype information is included.

The VEP can be used to analyze data from any species with an assembled genome sequence and an annotated gene set. The data files necessary for annotation in 80 vertebrate species and many invertebrates are distributed by Ensembl and Ensembl Genomes (Kersey et al. 2014), respectively. These are updated regularly ensuring analysis can be performed using contemporary biological knowledge. The VEP also supports both the latest GRCh38 and previous GRCh37 human assemblies. Importantly, all results are fully reproducible using Ensembl archived versions. Finally, researchers may use their own transcript data for analysis, for example in species not yet in Ensembl, or for novel or private annotations. A script is included in the VEP script package to create an annotation set from a GFF and FASTA file pair.

Each version of the VEP is tied to a specific release of Ensembl. This explicit versioning ensures all results are stable across a release, which is critical for provenance and reproducibility. To avoid misinterpretation of a variant based on a previous transcript or protein version, the output includes the identifier and version in HGVS coding descriptions. The VEP is open source, free to use, actively maintained and developed. A mailing list (http://lists.ensembl.org/mailman/listinfo/dev) provides responsive support and the benefits of a shared community. The wide usage helps ensure bugs are found and corrected rapidly, and enables suggestions to be gathered from a broad range of project teams.

The nature of the VEP results are described below along with input and output formats, the different interfaces and details on performance.

### Transcript annotation

The VEP results include a wide variety of gene and transcript related information (see Table 2). Any transcript set on a primary reference assembly or on ALT sequences can be used, but the VEP selects Ensembl annotation by default. For *Homo sapiens* and *Mus musculus* this is the GENCODE gene set, which denotes that it is a full merge of Ensembl’s evidence-based transcript predictions with manual annotation to create the most extensive set of transcripts isoforms for these species (Frankish et al. 2015). The Ensembl transcripts match the reference genome assembly exactly, which eliminates the potential for errors in annotation due to differences between the reference and transcript annotation. If configured to use the RefSeq transcript set, mismatches between a transcript and the genome reference assembly are reported to eliminate possible confusion in the interpretation.

**Table 2:** Gene and transcript-related fields reported by VEP

| **Property** | **Description** |
| --- | --- |
| Gene ID | Ensembl stable identifier for affected gene |
| Gene symbol | Common name for gene e.g. from HGNC |
| Transcript ID | Ensembl stable identifier for affected transcript |
| RefSeq ID | NCBI RefSeq identifier for affected transcript |
| CCDS ID | Consensus CDS identifier uniting Havana, Ensembl and NCBI |
| Biotype | GENCODE biotype of affected transcript |
| cDNA coordinates | Coordinates of input variant in unprocessed cDNA |
| CDS coordinates | Coordinates of input variant in processed coding sequence |
| Distance | Distance to transcript if variant falls outside transcript boundaries |
| Consequence type | SO consequence type of input variant allele on transcript |
| Exon | Number(s) of affected exon(s) |
| Intron | Number(s) of affected intron(s) |
| TSL | Transcript support level highlights well-supported and poorly-supported transcript models |
| APPRIS | APPRIS is a system to annotate alternatively spliced transcripts based on a range of computational methods, assigning primary and alternate statuses to transcripts |
| HGVS | HGVS notations for input variant relative to the coding sequence |
| Phenotype | Flag indicating known association with a phenotype or disease |

A variant may have more than one alternate non-reference allele and may overlap more than one transcript or regulatory region. Therefore, to present the most comprehensive annotation the VEP output reports one line (or unit) of annotation per variant alternate allele per genomic feature. As yet, there is no robust annotation of dominant transcript per tissue type available so the VEP includes a variety of data to help filter the many different transcript isoforms. For example, in *Homo sapiens* and *Mus musculus* the filtered GENCODE Basic transcript set includes the vast majority of transcripts identified as dominantly expressed (Frankish et al. 2015) and consensus coding sequence (CCDS) annotation highlights transcripts having the same CDS in both RefSeq and Ensembl. In several species, a ranking of supporting evidence for transcripts using Transcript Supporting Level data can prioritize consequences for review (TSL: http://www.ensembl.org/Help/Glossary?id=492), while APPRIS provides automated annotation of principal transcript isoforms (Rodriguez et al. 2013). Cross-references to known proteins in UniProt and the option to filter for variants in protein coding transcripts are also included. In *Homo sapiens*, for clinically relevant loci requiring stable annotation, the VEP can annotate on Locus Reference Genomic (LRG) sequences. Furthermore, the VEP has a flexible “plugin” architecture (described below) to enable for algorithmic extensions for additional analysis. For example an experimental plugin, GXA.pm, uses data from the Expression Atlas project (Petryszak et al. 2014) to indicate expression levels across tissues for many transcripts, which can be used to filter transcript isoforms.

### Protein annotation

Protein sequence changes are annotated with the information in Table 3. The VEP also provides an indication of the effect of the amino acid change using protein biophysical properties. These data can improve interpretation of protein variants with no associated phenotype or disease data by predicting how deleterious a given mutation may be on the functional status of the resultant protein. Scores and predictions are pre-calculated for all possible amino acid substitutions, and updated when necessary, ensuring that even the annotation of novel variants is rapid. Sorting Intolerant From Tolerant (SIFT, Kumar et al. 2009) results are available for the ten species that are most used in Ensembl. PolyPhen-2 (Adzhubei et al. 2010) results are available for human proteins. Other pathogenicity predictor scores such as Condel (Gonzalez-Perez et al. 2012), FATHMM (Shihab et al. 2013) and MutationTaster (Schwarz et al. 2014) are available for human data via VEP plugins (see Table 4).

**Table 3:**
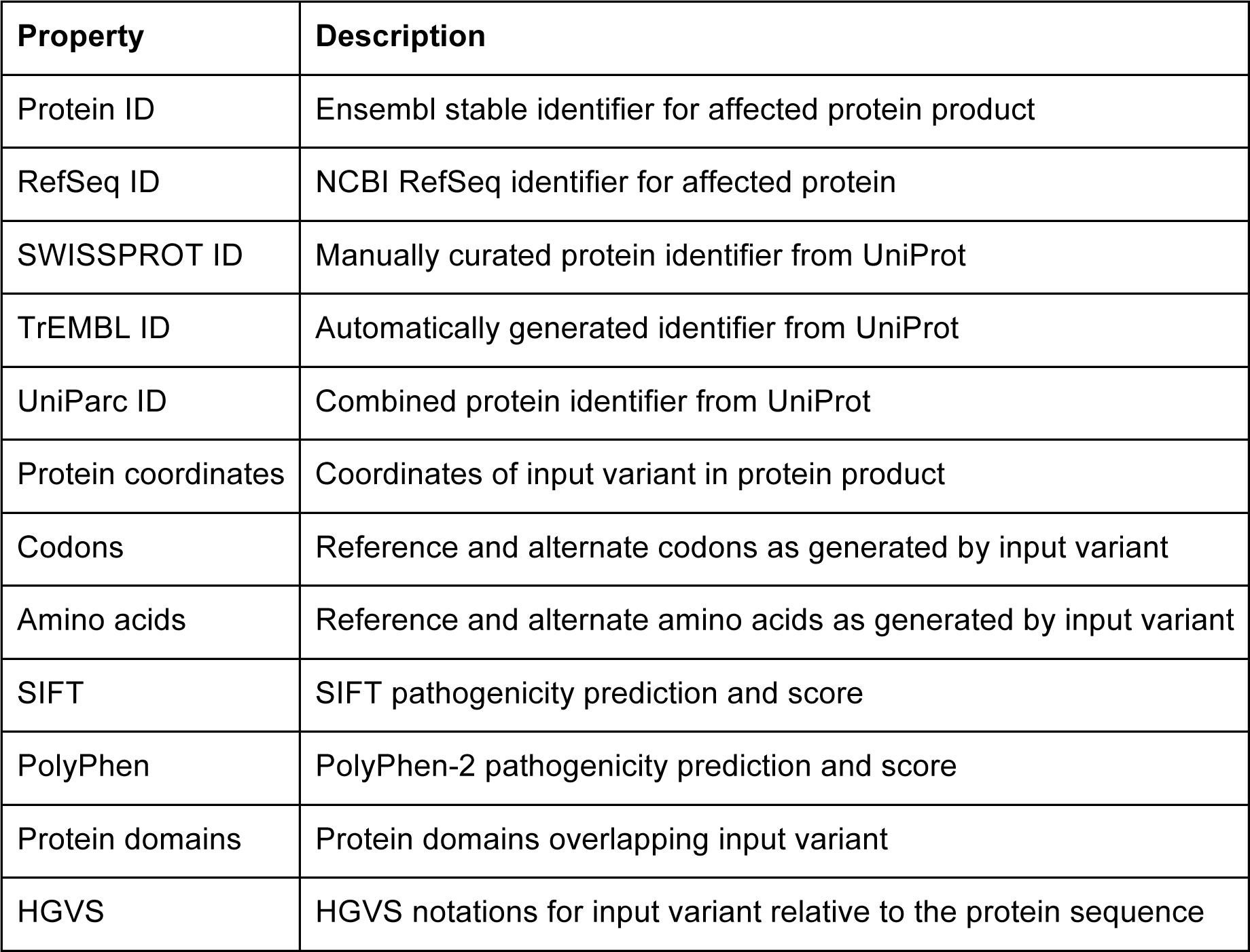
Protein-related fields reported by VEP

| Property | Description |
| --- | --- |
| Protein ID | Ensembl stable identifier for affected protein product |
| RefSeq ID | NCBI RefSeq identifier for affected protein |
| SWISSPROT ID | Manually curated protein identifier from UniProt |
| TrEMBL ID | Automatically generated identifier from UniProt |
| UniParc ID | Combined protein identifier from UniProt |
| Protein coordinates | Coordinates of input variant in protein product |
| Codons | Reference and alternate codons as generated by input variant |
| Amino acids | Reference and alternate amino acids as generated by input variant |
| SIFT | SIFT pathogenicity prediction and score |
| PolyPhen | PolyPhen-2 pathogenicity prediction and score |
| Protein domains | Protein domains overlapping input variant |
| HGVS | HGVS notations for input variant relative to the protein sequence |

**Table 4:** Examples of VEP plugins. For the full list see here: https://github.com/ensembl-variation/VEP_plugins

| Plugin | Maintained by | Functionality |
| --- | --- | --- |
| CADD | Martin Kircher | Integrates multiple annotations into one metric by contrasting variants that survived natural selection with simulated mutations |
| dbNSFP | Ensembl | Provides pre-calculated scores from dbNSFP for many pathogenicity prediction tools for every possible missense variant in the human genome (Liu et al. 2013). |
| dbscSNV | Ensembl | Retrieves data for splicing variants from dbscSNV (Jian et al. 2014) |
| ExAC | Ensembl | Retrieves ExAC allele frequencies from the Exome Aggregation Consortium (ExAC) project (http://exac.broadinstitute.org). |
| GWAVA | Graham Ritchie | Predicts the functional impact of variants on non-coding elements from e.g. ENCODE using GWAVA |
| GXA | Ensembl | Reports data from the Expression Atlas |
| LD | Ensembl | Finds variants in linkage disequilibrium with any overlapping existing variants |
| LOFTEE | Konrad Karczewski | Predicts if stop gain, splice site or frameshift variants lead to loss of function (LoF) in the affected protein |
| MaxEntScan | Ensembl | Compares scores for reference and mutant splice site sequences using a maximum entropy method |
| miRNA | Ensembl | Reports whether a variant is predicted to fall in a stem or loop region of a mature miRNA |
| UpDownStream | Ensembl | By default the VEP searches 5 kb either side of input variants for transcripts. Configures this distance which is useful in species with small intergenic distances, or for investigating long range trans-acting regulatory interactions |
| VAX | Michael Yourshaw | Incorporates data from KEGG, Human Protein Atlas, MitoCarta, OMIM and more into VEP output |

### Non-coding annotation

Variants in non-coding regions may have an impact on transcriptional or translational regulation if they fall in regulatory regions. The VEP reports variants in non-coding RNAs, genomic regulatory regions or transcription factor binding motifs, and also reports changes to the consensus score of binding motifs (see Table 5), which have been shown to be implicated in disease (Ward and Kellis 2012). The Ensembl Regulatory Build (Zerbino et al. 2015), which uses data from ENCODE (The ENCODE Project Consortium 2012), BLUEPRINT (Adams et al. 2012) and the NIH Epigenomics Roadmap (Romanoski et al. 2015), is the primary regulatory annotation but the VEP analysis can be limited to regulatory regions observed in specific cell types. GERP (Cooper et al. 2005) and other conservation scores derived from genomic multiple alignments, which may predict functional importance in non-coding regions, can be added via a plugin. GWAVA (Ritchie et al. 2014), CADD (Kircher et al. 2014) and FATHMM-MKL (Shihab et al. 2014) plugins are also available, which integrate genomic and epigenomic factors to grade and prioritize non-coding variants.

**Table 5:** Regulatory element-related fields reported by VEP

| Property | Description |
| --- | --- |
| Regulatory or Motif feature ID | Ensembl identifier for affected regulatory element |
| Motif name | External name for transcription factor binding motif |
| Motif position | Coordinates of input variant in transcription factor binding motifs |
| Motif score | Score reflecting effect of input variant on closeness of binding motif sequence to consensus |
| Informative position | Flag indicating if the position occupied by the variant in the binding motif is important in the consensus sequence |

### Frequency, phenotype and citation annotation

The VEP searches the Ensembl Variation databases, which contain a large catalogue of freely available germ line and somatic variation data in vertebrates (Chen et al. 2010; Rios et al. 2010). Ensembl integrates and quality checks variants from dbSNP (Sherry et al. 2001) and other sources for twenty species. Additional human data include mutations from COSMIC (Forbes et al. 2011) and the Human Gene Mutation Database (Stenson et al. 2012), and structural variants and copy number variants (CNVs) from the DGVa (Lappalainen et al. 2013). Therefore, the VEP can reference millions of variants to identify those previously reported. The VEP reports allele frequencies from the 1000 Genomes, NHLBI exome sequencing (http://evs.gs.washington.edu/EVS/) and ExAC projects. These can be used as filters, allowing common variants to be excluded as candidates for pathogenicity (see Table 6 for a list of the annotations provided). The VEP includes PubMed identifiers for variants which have been cited, and also annotates those associated with a phenotype, disease or trait, using data from OMIM (http://omim.org/), Orphanet (http://www.orpha.net/), the GWAS Catalog (Welter et al. 2013) and other data sources (http://www.ensembl.org/info/genome/variation/sources_phenotype_documentation.html). Clinical significance states assigned by ClinVar (Landrum et al. 2014) are also available for human variants.

**Table 6:** Co-located variant-related fields reported by VEP

| Property | Description |
| --- | --- |
| Variant ID | External identifier for variant co-located with input e.g. rsID from dbSNP |
| Somatic | Somatic status of co-located variant |
| GMAF | Global minor allele and frequency of co-located variant from combined 1000 Genomes phase 3 populations |
| Other frequencies | Frequency data from continental level 1000 Genomes phase 3 data and two NHLBI-Exome Sequencing Project populations |
| Clinical significance | Clinical significance status of co-located variant as reported by ClinVar |
| Phenotype | Flag indicating known association with a phenotype or disease |
| PubMed ID | NCBI PubMed IDs of publications citing co-located variant |

### Input and output formats

The VEP supports input data in variant call format (VCF), the standard exchange format used in NGS pipelines. Unlike other tools (see Table 1), the VEP can also process variant identifiers (e.g. from dbSNP) and HGVS nomenclature notations (e.g. HGVS using Ensembl, RefSeq or LRG transcripts and proteins ‘ENST00000615779.4:c.102944T>C’; ‘BRCA2:p.Val2466Ala’; ‘Q15118:p.Val42Phe’). These identifiers are commonly used in publications and reports. This functionality can also be used to “reverse map” variants from cDNA or protein coordinates to the genome and vice versa.

VEP output consists of an HTML or text format summary file and a primary results file in tab-delimited, VCF, GVF or JSON format. The default tab-delimited output is designed to present key data in a human-readable format that is easily parsed and can include detailed and complex data alongside. The VEP’s VCF output follows a standard agreed with other annotation tool providers (Cingolani et al. 2012) to promote transparent cross-comparison and benchmarking of results.

Variant consequences are described using a standardized set of variant annotation terms (http://www.ensembl.org/info/genome/variation/predicted_data.html#consequences) which were defined in collaboration with the Sequence Ontology (SO) (Cunningham et al. 2015b). Each consequence term has a stable identifier and definition, thereby removing ambiguity in definition or meaning. Structuring the consequences ontologically enables powerful querying: it is possible to retrieve all coding variants in one query without the need to specify each sub-category such as stop_gained, missense, synonymous etc. The SO terms are used widely including by the UCSC Genome Browser (Rosenbloom et al. 2014), the 1000 Genomes Project (Clarke et al. 2012), ClinVar, the ExAC project and the International Cancer Genome Consortium (The International Cancer Genome Consortium Mutation Pathways and Consequences Subgroup of the Bioinformatics Analyses Working Group 2013) allowing transparent interoperability and cross-validation.

### VEP interfaces

The VEP is platform independent and available as (1) an online tool, (2) an easily installed Perl script, or (3) via the Ensembl Representational State Transfer (REST) application program interface (API) (Yates et al. 2014). Each interface is optimized to support different quantities of data and levels of bioinformatics experience. All three use the same core codebase to ensure results are consistent across each interface. A comprehensive test suite backs all code, with continuous integration performed by Travis CI (https://travis-ci.org/) ensuring high quality code, which must pass stringent quality tests before release.

#### 1) VEP Web

VEP Web (http://www.ensembl.org/vep) offers a simple point-and-click interface. This is ideal for exploring annotation in an interactive manner. The portal is most suited to first-time use or small-scale analysis. The maximum compressed uploaded data file size currently supported is 50 megabytes, large enough for around 2 million typical lines of VCF data.

For single variant analysis, the web interface incorporates ‘Instant VEP’ functionality. Pasting or typing a single variant such as a variant in HGVS notation from a manuscript will rapidly return basic consequence prediction data. To submit a request for more than one variant, data can be uploaded, pasted or given via URL and options selected using a simple online form. A limited set of the VEP’s most commonly used plugins is available to use via the web interface. Requests are processed by a resource management system on the Ensembl web servers to distribute the request load.

The output web page (see example in Figure 2) shows summary statistics and charts to provide an overview of the results. It also has a table with a preview of the detailed results, with a simple interface to configure filtering of the output. Via a series of drop-down menus, multiple filters (see examples in Table 7) can be combined using basic logical relationships thereby allowing the creation of complex customized queries. This is designed to aid prioritization of smaller numbers of variants. Results can be stored by logging into an Ensembl account.

**Figure 2:**
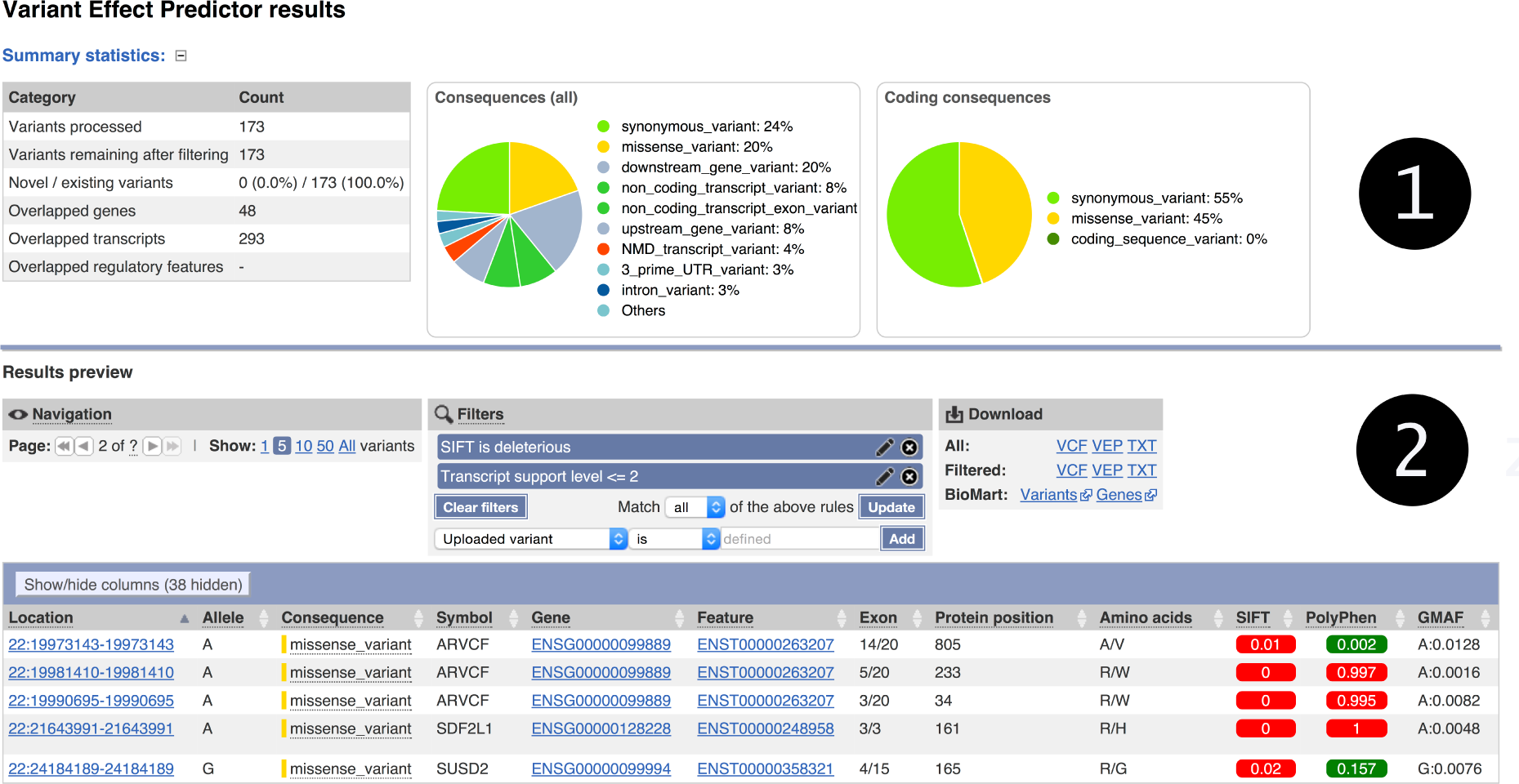
A typical VEP Web results page. Section (1) gives summary pie charts and statistics. Section (2) contains a preview of the results table with navigation, filtering and download options. The preview table contains hyperlinks to genes, transcripts, regulatory features and variants in the Ensembl browser. The results can be downloaded in VCF, text or a custom VEP file formats.

**Table 7:** Example filters available in VEP

| Option or command | Description |
| --- | --- |
| **Runtime filters** |  |
| --no_intergenic | Filter out variants that fall in intergenic regions |
| --pick | Choose one consequence for each variant; priority is given to the canonical transcript for each gene, protein coding transcripts, and more severe consequence types e.g. missense_variant is more severe than intron_variant |
| --per_gene | Picks one consequence using the same methodology as --pick but chooses one per overlapping gene |
| --filter_common | Filter out variants that are co-located with a known variant that has a minor allele frequency greater than 1%. |
| **Results filters using filter_vep.pl** |  |
| SIFT is deleterious OR PolyPhen is probably_damaging | Filter for results where SIFT or PolyPhen-2 predicts the variant protein will be non-functional |
| AFR > 0.1 AND EUR < 0.05 | Filter for variants co-located with those that are common in African populations but rare in European populations |
| Gene in gene_list.txt AND Phenotype matches cancer | Filter for results for variants that fall in the genes with IDs listed in gene_list.txt and that have been annotated with a cancer phenotype from a custom dataset (VEP script only) |

#### 2) VEP Script

The downloadable Perl script (http://www.ensembl.org/info/docs/tools/vep/script/index.html) is the most powerful and flexible way to use the VEP. It supports more options than the other interfaces, has no limit on input file size and includes extensive input, output, filtering and analysis options.

To install the script, simply download the VEP package and run the installer script, which automatically downloads the necessary API and annotation files (or ‘cache’ files). Updates with the latest data are available for each Ensembl release. The full source code is freely available on the Ensembl GitHub repository.

To process large volumes of data, the VEP script works most efficiently in ‘offline’ mode using a local cache of transcript annotations rather than online public databases. As well as optimizing runtime, this ensures data privacy for clinically or commercially sensitive data. Furthermore, the VEP input can be configured to query overlaps with local, potentially private, variant and phenotype data or other custom data sets. In this manner annotation in formats including BED, GFF, GTF, VCF and bigWig formats can be incorporated into the VEP output.

Advanced filtering options are available for a smaller result set, either during runtime or as a post-run process, see Table 7. Filtering can be performed as a post-run process by an accompanying script that uses a simple field-operator-value language. Filtered results can be fed back to the VEP for further analysis or exported.

With some familiarity of Perl, the VEP can truly be customized, extended and integrated with other systems. As almost all of the algorithmic content of the VEP is contained within the Ensembl API, the features of the VEP can be accessed using API calls. It is therefore trivial to extend the VEP results and perform secondary analyses, such as retrieving all OMIM IDs associated with the genes in the VEP results, or calculating known variants in linkage disequilibrium with a subset of variants. Alternatively, the VEP is also customizable via its plugin architecture, which was developed to provide greater scope for expansion. This architecture supports the use of VEP as the backbone of a customized analysis pipeline by writing additional code to extend the VEP’s functionality for specific use cases. Example uses include filtering output, adding annotation from local or remote sources, executing external programs or rendering graphical representations of the output. Ensembl provides a number of VEP plugins, hosted on GitHub (https://github.com/Ensembl/VEP_plugins), and a variety are published (Ritchie et al. 2014; Yourshaw et al. 2014) (see Table 4).

#### 3) VEP REST API

Ensembl’s language-independent REST API provides robust computational access in any programming language and returns basic variant annotation and consequence data. Individually or in batches of up to 1000, variants can be submitted to the API server in a single request. Results return in JSON, simple for parsing in most modern programming languages (see Figure 3 for an example of JSON output). Using this interface dynamic VEP queries can be integrated into custom-built software for on-demand results, as used for example in the Decipher Genome Browser (Bragin et al. 2014). For documentation see: http://rest.ensembl.org/#VEP.

**Figure 3:**
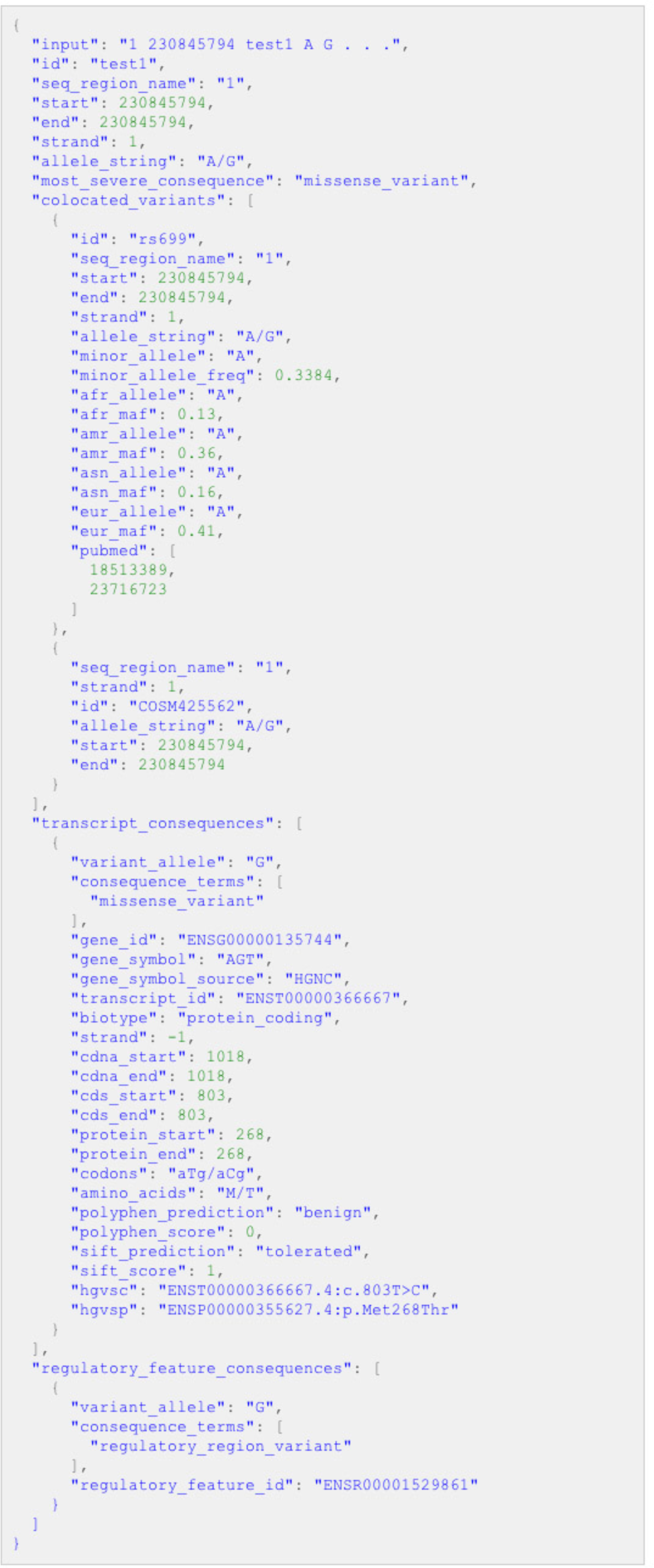
Example of JSON output as produced by the VEP script and REST API (redacted and prettified for display).

As with the web interface, a limited set of the VEP’s most commonly used plugins is configured for use via the REST API.

### Performance

The VEP script can be threaded for rapid performance on systems with multiple CPU cores. A typical human individual’s variant set can be processed in around an hour on a modern quad core machine; the 4,474,140 variants in NA12878 from Illumina’s Platinum Genomes set (ftp://ussd-ftp.illumina.com/hg19/2.0.1/NA12878/) took 62 minutes to process (see Table 8). This reduces to 32 minutes using the smaller GENCODE basic gene set. A negligible startup time means the VEP achieves similar throughput rates on both small and large datasets. A typical exome sequencing data set (100,000 to 200,000 variants) is processed in under five minutes.

**Table 8:**
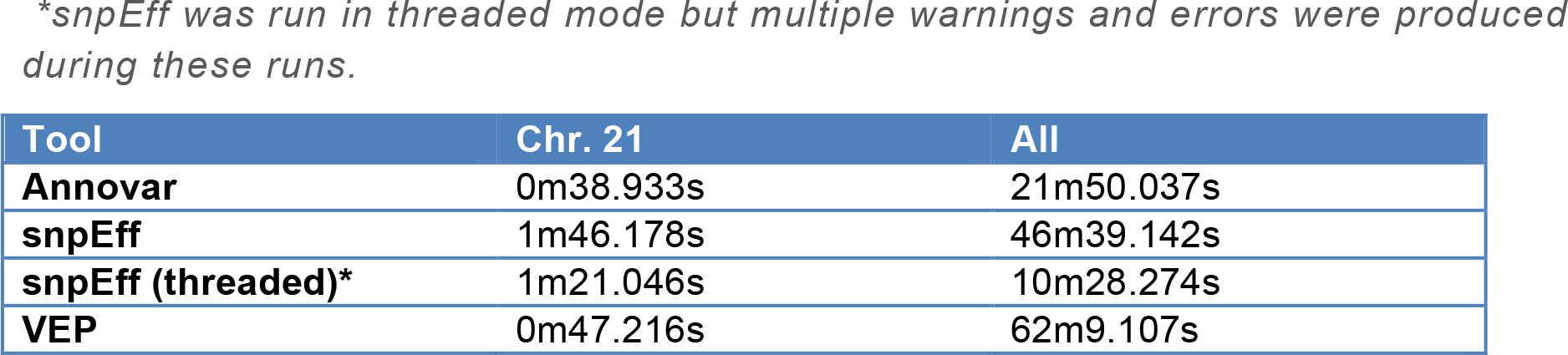
Comparison of runtime. Two datasets from Illumina’s Platinum Genomes were used (http://www.illumina.com/platinumgenomes/), both on the GRCh37 assembly: 67416 variants from chromosome 21, and the whole genome set of 4474140 variants. Each tool was configured to use the Ensembl release 75 gene set, with options configured for the fastest runtime. ^*^snpEff was run in threaded mode but multiple warnings and errors were produced during these runs.

| Tool | Chr. 21 | All |
| --- | --- | --- |
| **AnnoVar** | 0m38.933s | 21m50.037s |
| **snpEff** | 1m46.178s | 46m39.142s |
| **snpEff (threaded)\*** | 1m21.046s | 10m28.274s |
| **VEP** | 0m47.216s | 62m9.107s |

To improve runtime, individual VEP jobs can be threaded across multiple processor cores. Larger scale parallel processing architectures such as compute farms enable further subdivision of the VEP job (for example, by chromosome).

The VEP’s runtime performance is compared to Annovar and snpEff in Table 8. For smaller input files, the VEP performs as well as or faster than other tools. The VEP concedes time to snpEff by being written in Perl (an interpreted language) versus compiled Java for snpEff (http://www.codeproject.com/Articles/696764/Differences-between-compiled-and-Interpreted-Langu). Annovar, while also written in Perl, does not provide the same depth of annotation as VEP and so runs faster. It should also be noted that the VEP, through the REST API or through the Instant VEP functionality of the VEP web interface, returns predictions for single variants in a fraction of a second. This is available to users without any software download or installation, something neither Annovar nor snpEff can offer.

Run time varies with the number and complexity of overlapping genomic features, resulting in faster analysis times for species with sparse annotation than those with rich annotation such as human and mouse.

As the web and REST implementations are based on the same underlying code as the VEP script, performance is broadly comparable to the above, with allowances made for job queues (for web), network transfer of data (for web and REST) and request limits (for REST).

## DISCUSSION

The Ensembl Variant Effect Predictor software provides tools and methods for a systematic approach to annotate and prioritize variants in both large scale sequencing projects and smaller analysis studies. By automating annotation in a standard manner and reducing the time required for manual review, it helps manage many of the common challenges associated with analysis of SNPs, short insertions-deletions, copy number variants and structural variants. The VEP annotates variants using a wide range of reference data including: transcripts, regulatory regions, frequencies from previously observed variants, citations, clinical significance information and predictions of biophysical consequences of variants.

The quality, quantity and stability of variant annotation obtained depends on the choice of transcript set used (McCarthy et al. 2014). As such, the VEP allows flexibility of transcript choice. To effectively manage large numbers of variant annotations and transcript isoforms, the VEP provides several methods to prioritize results and reduce the number of variants needing manual review. A selection of these filters is available, but in addition, the VEP supports building of custom filters. Uniquely, the VEP algorithm can be expanded to perform additional calculations via plugins (as exemplified in Yourshaw et al. 2014) and can analyze custom, potentially private, data.

Interpreting all variants in a genome remains an unsolved challenge. An increasing number of large-scale WGS will detect rare variants in both coding and non-coding regions of the genome and further possible identification of loci associated with disease. Having these variants available in public repositories such as dbSNP and the European Variant Archive, or discoverable using federated resources will be of significant benefit for analysis. Emerging efforts such as the Global Alliance for Genomic Health (GA4GH) Beacon project (https://beacon-network.org/) are currently developing possible distributed solutions.

Improved functional annotation is especially critical for variants in non-coding regions. Many fall in loci that regulate gene expression in specific tissues. Characterizing associations between transcripts and tissues will facilitate a subset of tissue-specific transcript isoforms to be selected for variant annotation, tailoring results. Moreover, upon providing the link from regulatory region to regulated gene, the potential molecular mechanism underlying disease could be explained. Data from large scale efforts such as the GTEx project, which aims to systematically characterize the effects of regulatory variants in different tissues (GTEx Consortium et al. 2015), will be integrated into the VEP reference data in order to have the most current data available to the VEP for analysis.

As discussed above, standardized SO terms are used for describing variant consequences and VEP results can be output in VCF format. Work is ongoing to develop a comprehensive variant annotation data exchange format within the GA4GH. Furthermore, the GA4GH is defining standards for representation of associations between variants and phenotypes, traits and diseases. The VEP will support such formats when they are mature.

The VEP is also regularly extended and improved (see release notes http://www.ensembl.org/info/docs/tools/vep/script/vep_download.html#history) with new features added to both the core VEP code and the plugin library. To improve genome-wide analysis, the VEP will leverage data from future sequencing projects, implement new algorithms and adopt data exchange standards and, therefore bring continual benefit to variant interpretation.

## METHODS

The VEP algorithms and code are part of the freely available Ensembl API, coded in the Perl programming language. Time-critical components are written in C and incorporated into the API using the XS framework (http://perldoc.perl.org/perlxs.html). Installation of the VEP script triggers automated installation of the Ensembl API, along with the BioPerl API (Stajich et al. 2002) upon which the Ensembl API depends. All interfaces to the VEP use the same underlying API calls, ensuring consistency across the different VEP access platforms when version control is observed.

To process the input data, sequential contiguous blocks of variants (default block size 5000) are read into an input memory buffer. Each variant is converted into an Ensembl VariationFeature object that represents a genomic location and a set of alleles. Variants in tab-delimited and Pileup formats are converted directly to objects; those in HGVS notation are resolved to their genomic coordinates by extracting the relevant reference feature (transcript, protein or chromosome) using the Ensembl API. VCF input undergoes preprocessing to account for differences in how VCF and Ensembl represent unbalanced substitutions and indels. Optionally the parser can be forced to reduce each alternate allele to its most minimal representation by stripping identical bases from the 5′ and 3′ ends of the VCF reference and alternate allele in a process analogous to that described in (Tan et al. 2015). When using VEP’s forking functionality, the input buffer is divided amongst a number of sub-processes. Each sub-process carries out the analysis described hence, and then the results are rejoined and sorted back into input order before being written to output.

Input variants pass through a configurable quality-control process that checks for irregularities and inconsistencies. Variants that fail are reported via standard error output and/or in a warnings file. Checks include, for example, that allele lengths match input coordinates and the input reference allele matches that recorded in the reference genome.

The genomic loci overlapped by the variants in the input buffer are resolved to distinct megabase-sized regions. Each region corresponds to a single file on disk in the VEP cache, which contains objects serialized using Perl’s Storable framework (http://perldoc.perl.org/Storable.html). For each region, the transcripts, regulatory features and known variants are loaded from disk, deserialized into objects and cached in memory. This avoids rereading from disk when the same region is overlapped by variants in consecutive input buffers. The publicly available Ensembl databases can be used in place of the cache files to avoid downloading the data in advance, though doing so incurs a performance penalty due to network transfer rates.

Transcripts have a configurable flank (default 5000 base pairs) to allow the VEP to assign upstream and downstream status to variants within the region flanking a transcript. A hash-based tree structure is used to search for overlaps between input variants and genomic features. For each overlap, a VariationFeatureOverlap object is created, with specific subclasses for each genomic feature type: TranscriptVariation, RegulatoryFeatureVariation, MotifFeatureVariation. Each VariationFeatureOverlap object has two or more child VariationFeatureOverlapAllele objects representing each allele of the input variant – one representing the reference allele and one or more representing each of the alternate or mutant alleles. These objects are also sub-classed, with, for example, a TranscriptVariationAllele representing one allele of a variant overlapping a Transcript object.

For each TranscriptVariationAllele object, the API evaluates consequence types using a set of predicate functions. These assess whether, for example, a variant is predicted to cause a change in protein coding sequence (e.g. missense_variant). Prior to this, a series of pre-predicate checks are performed to improve runtime; for example, a variant does not need to be assessed for change to the protein sequence if it falls entirely within the intron of a transcript. These pre-predicate checks are also cached at each object “level”; for example, the location of a variant relative to the transcript structure is fixed at the TranscriptVariation level, but the allele type can be different for each TranscriptVariationAllele. The pre-predicate checks improve runtime by a factor of around two on a typical resequencing-based input file. Without them, runtime is proportional to *nfp*, where n is the number of input variant alleles, *f* is the number of overlapped features and *p* is the number of predicates; depending on a number of factors this can become as low as *nfp* /2 with pre-predicate checks enabled.

Predicates also make extensive use of caching: UTR, coding and translated sequences are all cached on the Transcript object with intron structure and other frequently accessed data. Established components of the Ensembl API handle tasks such as splicing exons and retranslating mutated sequences. Alternate codon tables are used as appropriate for mitochondrial sequences and selenocysteines. If a predicate is true for a given TranscriptVariationAllele, an OverlapConsequence object is assigned representing the consequence type; this object contains the appropriate SO term along with synonyms and ranking information. Each OverlapConsequence object type corresponds to one predicate. Hierarchy in the predicate system preserves the tree structure of the SO such that only the most specific term that applies under any given parent term is assigned; this same tree structure allows for ontological-style querying and filtering of the results. Multiple OverlapConsequence objects may be added to a single VariationFeatureOverlapAllele or TranscriptVariationAllele object to allow for complex cases, such as a variant that falls in a splice-relevant region that also affects the coding sequence of the transcript.

HGVS notations are also derived from TranscriptVariationAlleles, though they undergo significant additional processing to conform to the nomenclature definition (den Dunnen and Antonarakis 2000). For example, insertions or deletions with respect to the transcript sequence must be reported at the most 3′ position possible when they fall in repetitive sequence.

VariationFeatureOverlapAllele objects are then converted for writing to output, a process that involves several extra stages. VariationFeatureOverlapAlleles can be filtered in various ways which can be configured, for example: reporting only one VariationFeatureOverlapAllele per input variant; removing intergenic VariationFeatureOverlapAlleles (i.e. those produced from variants that don’t overlap a genomic feature); filtering based on allele frequency of a co-located known variant. Additional data fields are retrieved at this stage from relevant objects, for example: external identifiers for transcripts (UniProt, CCDS); exon and intron numbers; clinical significance for co-located variants. It is also at this stage that any configured plugins are executed. They are passed the VariationFeatureOverlapAllele object, which has accessor methods for other objects e.g. the Transcript, VariationFeature or genomic Slice. As plugin modules are executed after the VEP consequence calculation they have access to the VEP and Ensembl API objects before output data is written and return a data structure that is incorporated alongside the VEP’s main output data structure. The output data structure is then written to disk as one of several formats (tab-delimited, VCF, GVF, JSON), with the fields for each data format configurable at runtime. Output files contain headers describing the format and content of data fields, as well as version information for resources used.

### Cache and sequence files

The VEP’s caches are built for each of Ensembl’s primary species (70 species as of Ensembl version 84); the files are updated in concert with Ensembl’s release cycle ensuring access to the latest annotation data. Cache files for all previous releases remain available on Ensembl’s FTP archive site (ftp://ftp.ensembl.org/pub/) to facilitate reproducibility. For 15 of these species there are three types of cache files: one with the Ensembl transcripts, a ‘refseq’ one with the RefSeq transcripts and a ‘merged’ one that contains both. Caches for both the latest GRCh38 and previous GRCh37 (hg19) human genome builds are maintained. The human GRCh38 cache file is around five gigabytes in size, including transcript, regulatory and variant annotations as well as pathogenicity algorithm predictions. Performance using the cache is substantially faster than using the database; analyzing a small VCF file of 175 variants takes five seconds using the cache versus 40 seconds using the public Ensembl variation database over a local network (performance can be expected to be slower when using a remote database connection).

The VEP can use FASTA format files of genomic sequence for sequence retrieval. This functionality is needed to generate HGVS notations and to quality check input variants against the reference genome. VEP uses either an htslib-based (http://www.htslib.org/) indexer or BioPerl’s FASTA DB interface to provide fast random access to a whole genome FASTA file. Sequence may alternately be retrieved from an Ensembl core database, with corresponding performance penalties.

Cache and FASTA files are automatically downloaded and set up using the VEP package’s installer script, which utilizes checksums to ensure the integrity of downloaded files. The installer script can also download plugins by consulting a registry. The VEP package also includes a script, gtf2vep.pl, to build custom cache files. This requires a local GTF or GFF file that describes transcript structures and a FASTA file of the genomic sequence.

## DATA ACCESS

Pre-built data sets are available for all Ensembl and Ensembl Genomes species from ftp://ftp.ensembl.org/pub/current_variation/VEP/. They can also be downloaded automatically during set up whilst installing the VEP: http://www.ensembl.org/info/docs/tools/vep/script/vep_download.html#download

## ACKNOWLEDGMENTS

John Peden from Illumina for modifications and improvements to the forking process. The Ensembl team for gene annotation, regulatory annotation, comparative annotation and user support. The VEP community who have helped to improve the VEP by giving feedback and bug reports on.

This work was funded by the Wellcome Trust (grant numbers WT095908 and WT098051) and the European Molecular Biology Laboratory. This work has also received funding from the European Union’s Seventh Framework Programme (FP7/2007-2013) under grant agreement number 200754 (GEN2PHEN) and under grant agreement number 222664 (Quantomics), and from the European Union’s Horizon 2020 research and innovation programme under grant agreement number 634143 (MedBioinformatics).

## AUTHOR CONTRIBUTIONS

FC, WM, SEH wrote the paper with contributions and guidance from PF. WM wrote the VEP with contributions from GR and AT. LG, SEH, WM and AT develop the underlying APIs and build the Ensembl Variation databases. HR and WM developed the web interface. FC and PF provided supervision.

## DISCLOSURE DECLARATION (INCLUDING ANY CONFLICTS OF INTEREST)

Paul Flicek is a member of the Scientific Advisory Board for Omicia, Inc.

